# RNF219 interacts with CCR4-NOT in regulating stem cell differentiation

**DOI:** 10.1101/2020.07.06.190728

**Authors:** Hao Du, Chen Chen, Yan Wang, Yang Yang, Zhuanzhuan Che, Xiaoxu Liu, Siyan Meng, Chenghao Guo, Manman Xu, Haitong Fang, Chengqi Lin, Zhuojuan Luo

**Author notes:** These authors contributed equally to this work (Hao Du, Chen Chen, Yan Wang). To whom correspondence should be addressed at: Zhuojuan Luo, PhD, School of Life Science and Technology, the Key Laboratory of Developmental Genes and Human Disease, Southeast University, Nanjing 210096, China, Chengqi Lin, PhD, School of Life Science and Technology, the Key Laboratory of Developmental Genes and Human Disease, Southeast University, Nanjing 210096, China.

## Abstract

Regulation of RNA stability plays a crucial role in gene expression control. Deadenylation is the initial rate-limiting step for the majority of RNA decay events. Here, we show that RING Finger Protein 219 (RNF219) interacts with the CCR4-NOT deadenylase complex. RNF219-CCR4-NOT exhibits deadenylation activity *in vitro*. RNA-seq analyses identify some of the 2-cell specific genes and the neuronal genes significantly down-regulated upon RNF219 knockdown, while up-regulated after depletion of the CCR4-NOT subunit CNOT10 in mouse embryonic stem (ES) cells. RNF219 depletion leads to impaired neuronal lineage commitment during ES cell differentiation. Our study suggests that RNF219 is a novel interacting partner of CCR4-NOT, and required for maintenance of ES cell pluripotency.

## INTRODUCTION

Regulation of RNA metabolism is essential in various biological processes. RNA decay is the last but critical step to control RNA in both quantity and quality. Nearly all eukaryotic messenger RNAs (mRNAs) and long non-coding RNAs (lncRNAs) are protected by 7-methylguanosine (m^7^G) cap at the 5’-end and poly-adenine (poly(A)) tail at the 3’-end from degradation by exonucleases. RNA decay is usually triggered by transcript deprotection, such as decapping or deadenylation. In eukaryotes, deadenylation, or poly(A) tail shortening and removal, is the preferred initiating event for the vast majority of mRNAs and lncRNAs to be fated to rapid decay. After the rate-limiting deadenylation, unprotected bodies of RNA can be attacked at both ends by the 5’-3’ or 3’-5’ degradation machineries. In addition to mRNA and lncRNA decay, deadenylation is also vital for translational silencing.

To date, three types of deadenylase have been identified in mammals, including the CCR4-NOT complex, the PAN2-PAN3 complex and PARN (1). PARN maintains the short (A) tails of mRNAs in oocytes to control translation in a dormant state (2), and also functions in the maturation of snoRNA (3). PAN2-PAN3 is inefficient in trimming the final 20-25 adenosines of the poly(A) tail, thus minimally degrading the body of transcript (4,5). However, PAN2-PAN3 is able to function together with CCR4-NOT, which is the major deadenylase, to remove the poly(A) tail thoroughly and stimulate RNA decay (6). CCR4-NOT is a highly conserved multi-subunit complex, which contains two deadenylases, CNOT7 (or its paralogue CNOT8) and CNOT6 (or its paralogue CNOT6L)^(7)^. These two catalytic components differ in substrates: CNOT7 prunes poly(A) tail free of poly(A) binding protein (PABP), while CNOT6 dislodges protective PABP from the A tails, subsequently leading to deadenylation (8,9). CNOT1, the largest component of CCR4-NOT, interacts with most of the other components and serves as a scaffold required for the complex assembly. The RING finger containing CNOT4 possesses ubiquitin ligase activity in both yeast and human. CNOT4 controls the proteasome dependent degradation of multiple chromatin related proteins, such as the transcription regulator PAF1 and H3K4me3 demethylase JARID1C (10,11). Although the precise roles played by CNOT4 and other non-enzymatic components within CCR4-NOT remain to be elucidated, a growing body of evidence manifests that these components are indispensable for regulation of gene expression (12–14).

CCR4-NOT can be recruited to different subclasses of RNAs via different mechanisms. CCR4-NOT was found to engage in the deadenylation of RNAs with MicroRNA (miRNA) target sites by the miRNA-induced silencing complex (miRISC) complex ^(15–17)^. N6-Methyladenosine (m^6^A) reader protein YTHDF2 recruits CCR4-NOT to deadenylate RNAs with m^6^A modifications (18). CCR4-NOT interacts with Tristetraprolin (TTP), which is an AU-rich element (ARE) containing RNA binding protein, to deadenylate RNAs with AREs (19,20). The SMG5-SMG7 heterodimer recruits CCR4-NOT, thus triggering non-sense mediated mRNA decay (NMD) to rapidly degrade aberrant mRNAs bearing premature translation termination codons (21). However, other than these recruiting partners, factors involved in deadenylation through interacting with CCR4-NOT remain largely unknown.

RING Finger Protein 219 (RNF219), containing an evolutionarily conserved RING finger domain at its N-terminus, is a poorly characterized ubiquitin ligase. A recent study shows that RNF219 is able to promote DNA replication origin firing through ubiquitination of the ORC complex (22). A genetic variant of the RNF219 gene was reported to be associated with the risk for Alzheimer’s disease (23). Here we biochemically identified RNF219 as an interacting factor of CCR4-NOT. RNA levels of some of the 2-cell specific genes and the neuronal genes undergo the opposite regulation by RNF219 and the CCR4-NOT subunit CNOT10 in mouse ES cells. While the main subunits of CCR4-NOT are essential in preserving ES cell identity (24,25), we found here that depletion of RNF219 in ES cells leads to impaired neuronal lineage commitment. Our study suggests RNF219 as a cofactor of CCR4-NOT to regulate transcript levels, and also provides a novel direction in studying the pathological processes of *RNF219* variants linked human diseases, such as Alzheimer’s disease.

## RESULTS

### Interaction of RNF219 with the CCR4-NOT complex

In order to investigate the function of RNF219, we sought to identify factors that interact with RNF219. First, a tetracycline inducible stable cell line expressing N-terminal FLAG tagged RNF219 was generated by using the HEK-293 Flp-In-TRex (FIT) system. FLAG affinity purification was performed in the Benzonase nuclease treated condition to avoid nucleic acids dependent protein-protein interactions. FLAG-RNF219 and control purifications were then resolved by SDS/PAGE prior to silver staining (Fig 1A). Analyses of independent purifications from both nuclear and cytoplasmic S100 extracts led to the identification of nearly all the subunits of CCR4-NOT, except CNOT4 (Fig 1B). In addition to functioning in translation inhibition and deadenylation in cytoplasm, CCR4-NOT is also involved in RNA synthesis in nucleus at different stages, such as modification of the chromatin template, regulation of RNA polymerase II processivity (10,12). Thus, it is expected that the interaction between RNF219 and CCR4-NOT can be detected in both cytoplasm and nucleus.

**Figure 1.**
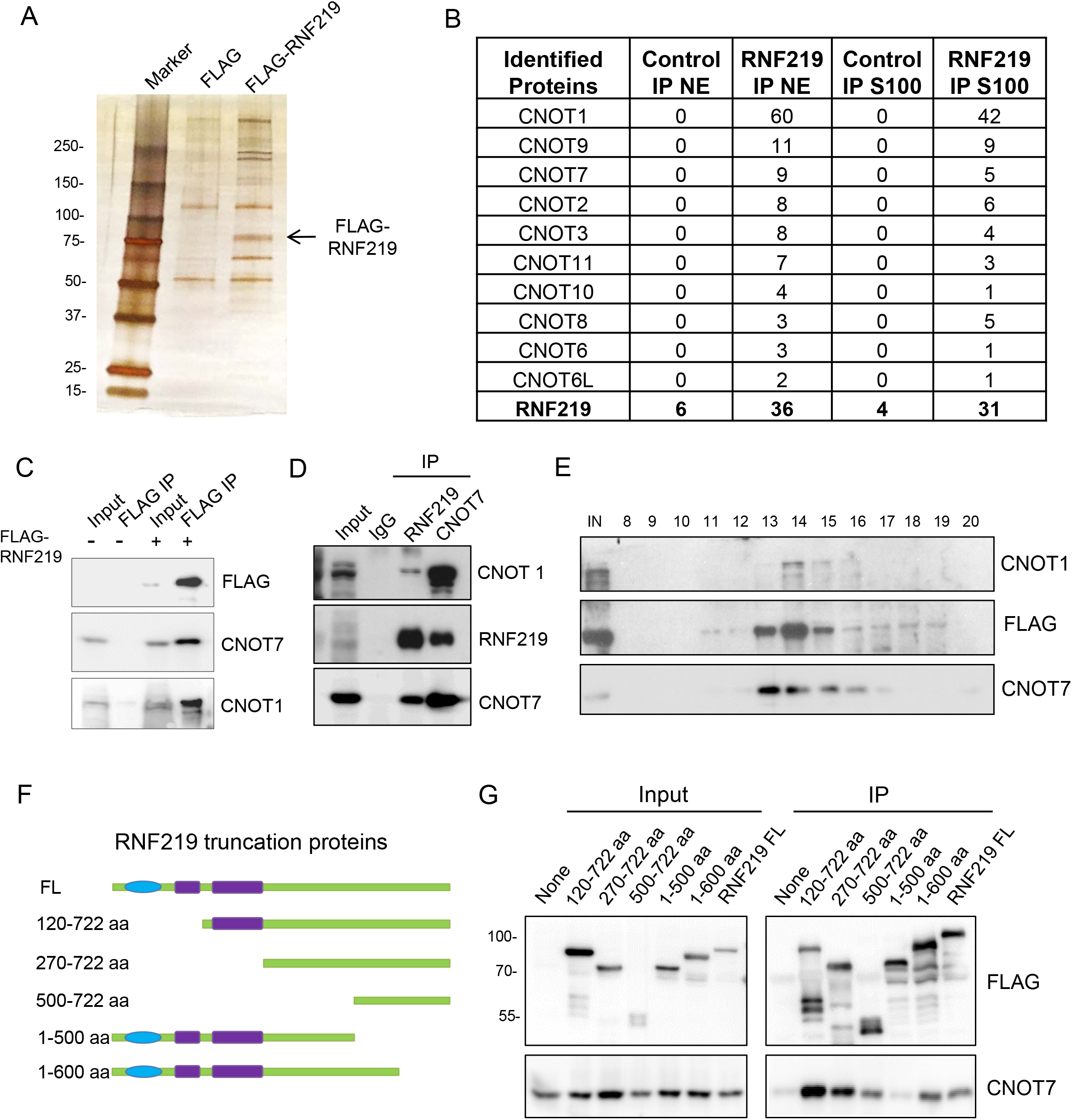
RNF219 interacts with the CCR4-NOT complex. (A) FLAG-purification of RNF219. Stable cell line expressing FLAG-tagged RNF219 was generated in HEK-293 FIT cells. The FLAG-RNF219 assoicated proteins were purified from the FLAG-RNF219 expressing cells using the FLAG-affinity purification method and analyzed by SDS-PAGE and silver staining. (B) Mass spectrometry identified the components of CCR4-NOT in the FLAG-RNF219 purification. S100: Cytoplasmic extract; NE: Nuclear extract. (C) Validation of the interaction of RNF219 with CNOT1 and CNOT7 by FLAG immunoprecipitation. (D) Endogenous co-IPs of RNF219 and CNOT7 from HEK-293 cells. (E) Size exclusion chromatography of cytoplasmic extracts from the FLAG-RNF219 stable cell line demonstrating that majority of RNF219 co-eluted with the components of CCR4-NOT at around 1.9 MDa (fractions 13–15). (F) Schematic diagram of RNF219 truncation proteins used in (G). (G) FLAG immunoprecipitations mapping the interaction domain of RNF219 with CNOT7. HEK-293 cells transfected with a plasmid encoding each of the FLAG-tagged RNF219 truncation protein (F) were subjected to cell lysis and FLAG immunoprecipitation.

The co-purification of RNF219 with the core components of CCR4-NOT, CNOT1 and CNOT7, were validated by Western blot (Fig 1C). Co-immunoprecipitation (co-IP) using the antibody specific to RNF219 or CNOT7 further confirmed the endogenous interaction between RNF219 and the CCR4-NOT components (Fig 1D; Fig S1A and S1B). In addition, co-fractionation of RNF219 with the CCR4-NOT components was examined by applying cytoplasmic and nuclear extracts from the HEK-293 FIT cells stably expressing FLAG-RNF219 to size exclusion chromatography. Western analyses revealed that the majority of RNF219 co-eluted with CNOT1 and CNOT7 at around 1.9 MDa (fraction 13-15; Fig 1E, S1C). The RNF219 protein consists of an N-terminal RING domain, a middle coiled-coil domain, and a C-terminal uncharacterized region (Fig 1F). A series of FLAG-tagged RNF219 truncated proteins were used to map the CCR4-NOT interaction domain in RNF219. No loss of binding to CNOT7 was observed when truncating both the RING and coil-coil domains (Fig 1G). The aa 1-600 C-terminal deletion mutant of RNF219 was able to interact with CNOT7, while the aa 1-500 mutant failed to co-immunoprecipitate CNOT7 (Fig 1G). Our results suggest that the aa 500-600 of RNF219 located in the center of the C-terminal region might mediate the interaction with CCR4-NOT.

### Deadenylation activity of RNF 219-CCR4-NOT

Give that CCR4-NOT is the major deadenylase in eukaryotic cells, we further explored whether RNF219 is also involved in deadenylation and RNA stability regulation. In order to examine whether RNF219 bound CCR4-NOT possesses deadenylation activity, we performed FLAG-CNOT1 and FLAG-RNF219 immuno-affinity purification from HEK-293 FIT cell lysates, respectively. The purified immune-precipitates were analyzed by Western blot and the catalytic subunit of CCR4-NOT, CNOT7, was co-purified with both FLAG-CNOT1 and FLAG-RNF219 (Fig 2A). *In vitro* deadenylation assay indicated that the affinity purified FLAG-RNF219 containing immuno-precipitate exhibited deadenylation activity toward the synthesized N20A20 RNA (Fig 2B).

**Figure 2.**
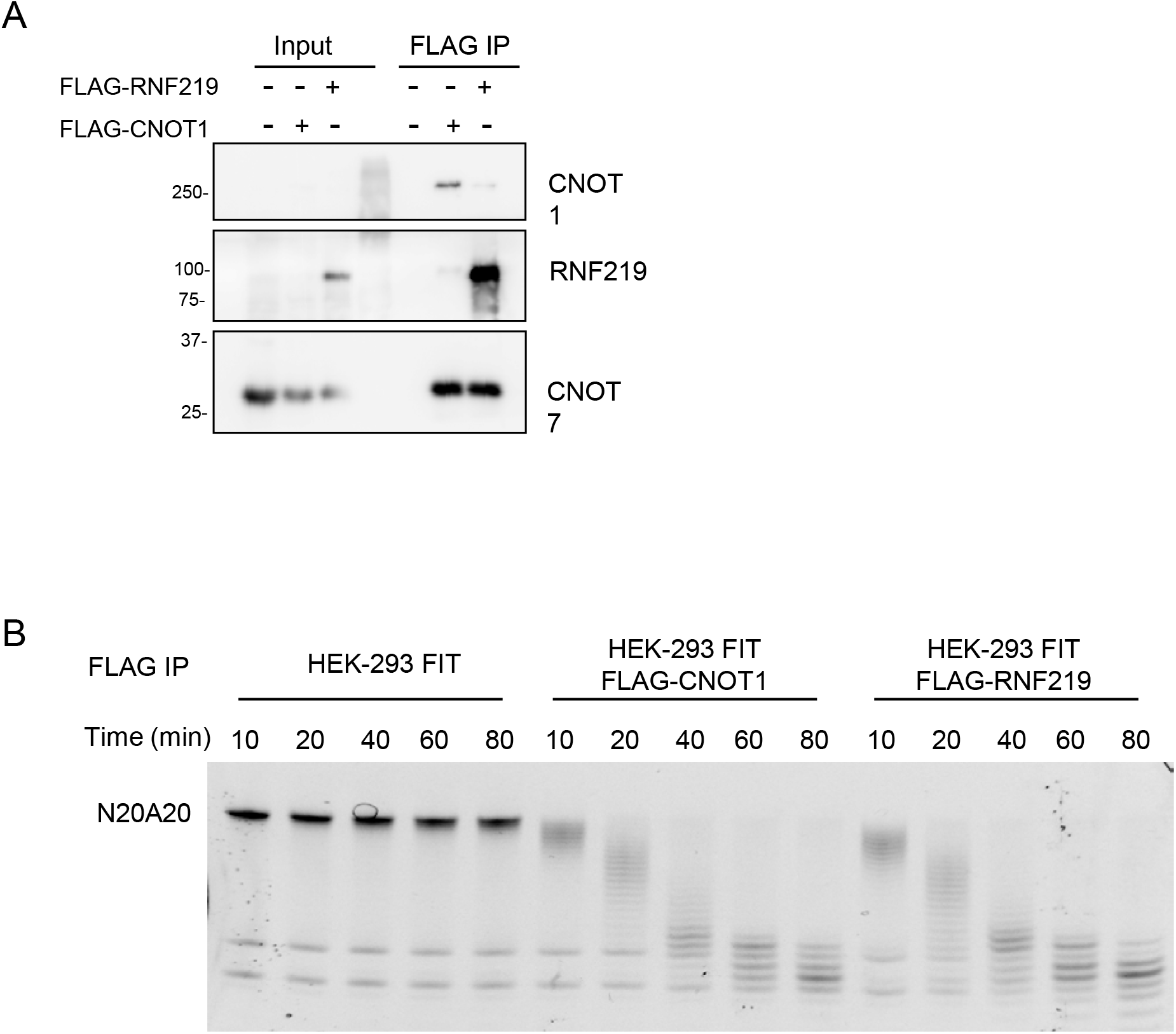
RNF219-CCR4-NOT exhibits deadenylation activity. (A) FLAG purification of RNF219 from the HEK-293 FIT cells stably expressing FLAG tagged RNF219, and CNOT1 from the FLAG-CNOT1 expressing HEK-293 FIT cells. Levels of RNF219, CNOT1 and CNOT7 in the purified immune-precipitates were examined by western blot. (B) *In vitro* deadenylation of the synthesized N20A20 RNA using the affinity purified FLAG-RNF219 or FLAG-CNOT1 containing immuno-precipitate. Samples were harvested at the indicated time intervals and resolved by electrophoresis separation. FLAG purification from HEK-293 FIT cell lysate was used as a negative control.

### Regulation of deadenylation by RNF219

To explore the potential mechanism underlying the role of RNF219 in maintaining transcript stability, we employed the transiently inducible b-globin (BG) reporter system combined with the transcriptional pulse-chase assay to analyze the effect of RNF219 on the deadenylation of RNA with different types of destabilization elements. As visualized by Northern blot, RNF219 overexpression significantly decelerated the deadenylation rate of the BG reporter that harbors the miRNA Let-7 target site (BG-L7) in the 3’ UTR (Fig 3B, S2A). As a control, RNF219 overexpression seemed not obviously affect the deadenylation of 3’ fragments of the basic BG reporter, which does not harbor any *cis*-element to promote deadenylation (Fig 3A). It has been previously established that the RISC complex can recruit CCR4-NOT to RNAs containing miRNA target sites. Here, we found that RNF219 was also able to interact with AGO2 in miRISC (Fig S2B). Therefore, it might be possible that RNF219 could modulate miRNA mediated deadenylation through physical interaction with CCR4-NOT and AGO2.

**Figure 3.**
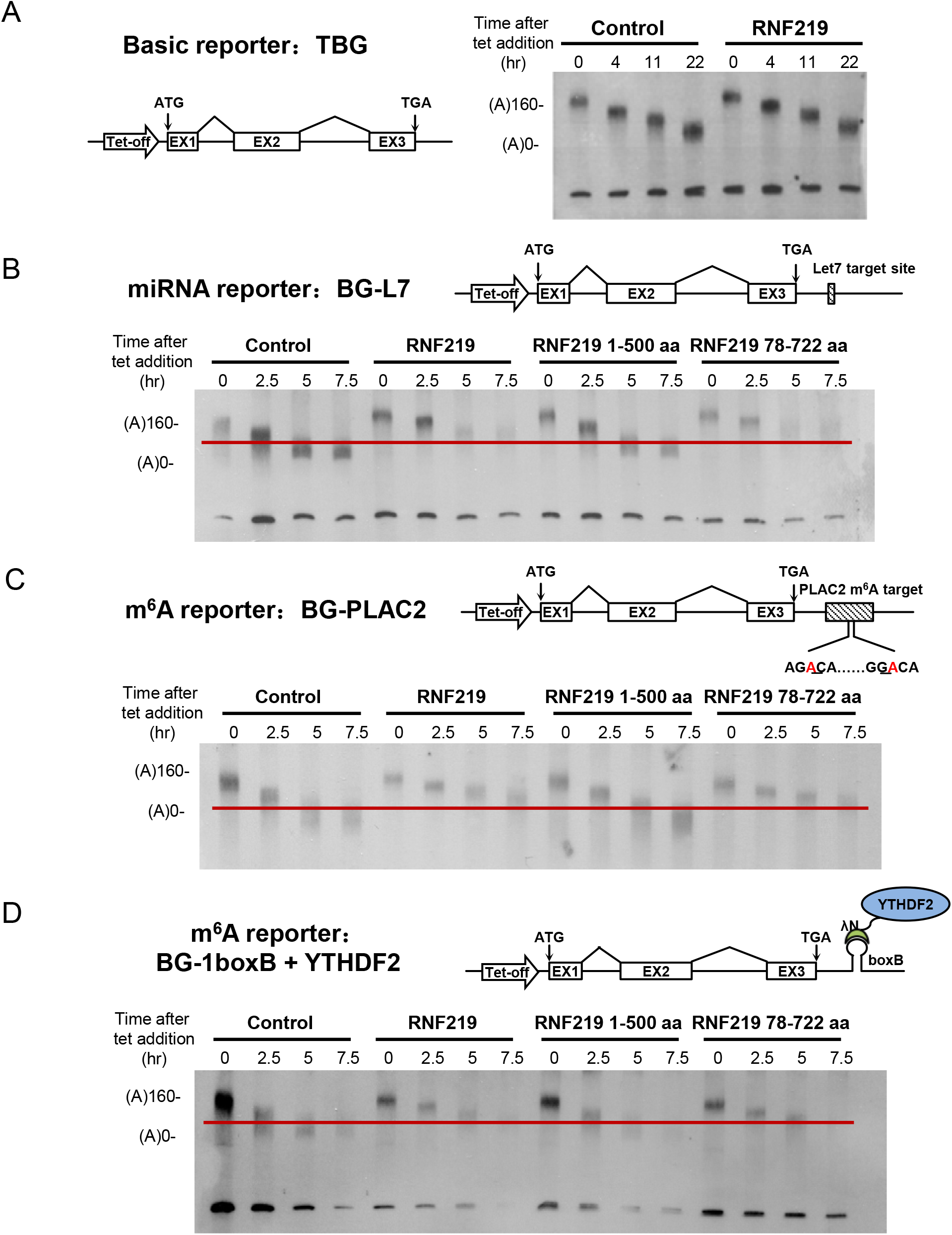
RNF219 affects deadenylation mediated by miRNA and m^6^A modification. (A) Schematic diagram of the basic reporter TBG construct. Blank boxes indicate exons (left panel). Deadenylation assay of TBG in control and RNF219 overexpressing cells (right panel). Tetracycline was briefly removed from the culture medium, thus leading to the production of a homogenous amount of BG mRNAs. Cells were harvested at the indicated time intervals for cytoplasmic RNA extraction. RNA samples were subjected to RNaseH cleavage, followed by electrophoresis separation and northern blot analyses. (B-D) Schematic diagrams of the miRNA reporter (B), the m^6^A reporter (C) and the YTHDF2 tethered reporter (D) constructs (upper panels). Deadenylation assay of each reporter following the overexpression of RNF219 full length, 1-500 aa and 78-722 aa (lower panels).

We next proceeded with the BG-PALC2 reporter, whose 3’ UTR contains a fragment with two m^6^A consensus motifs from the lncRNA PLAC2 (Fig 3C). Similar to its role in BG-L7, RNF219 also inhibited the deadenylation of the m^6^A reporter BG-PLAC2. YTHDF2, the m^6^A binding protein, is able to destabilize the m^6^A modified RNAs (28,29). To our expectation, RNF219 overexpression impeded the deadenylation of the YTHDF2 tethered BG reporter (Fig 3D). Thus, RNF219 might also regulate m^6^A mediated deadenylation.

To determine whether the role of RNF219 in the inhibition of deadenylation depends on its interaction with CCR4-NOT, we also carried out the deadenylation assay using the C-terminal deletion mutant of RNF219 (RNF219 1-500 aa), which lost the interaction with CCR4-NOT. As expected, RNF219 1-500 aa was incapable of slowing down the deadenylation rate of the transcripts of the BG-L7, BG-PLAC2 and YTHDF2 tethered reporters. In contrast, the RING domain deletion mutant of RNF219 (RNF219 78-722 aa), which retained the interaction with CCR4-NOT, was able to decelerate the deadenylation rate of these reporter transcripts to a similar extent as wild type RNF219 (Fig 3B-D). Thus, the inhibitory effect of RNF219 overexpression on deadenylation might require its interaction with CCR4-NOT.

### Opposite regulation of the 2-cell specific genes by RNF219 and CNOT10 in mouse ES cells

In order to further explore the functional link between RNF219 and the CCR4-NOT complex, we performed differential RNA level analyses using RNA-seq upon shRNA mediated knockdown of RNF219 or CNOT10, one of the core components of CCR4-NOT, in mouse ES cells. CNOT10 knockdown led to the RNA levels of 1,133 genes up-regulated, and 353 genes down-regulated (Fig S3A,B), consistent with the role of CCR4-NOT in negative regulation of RNA level by deadenylation. However, in total, only 489 genes were significantly differentially expressed after RNF219 knockdown, with 268 genes down-regulated (Fig 4A,B). These genes down-regulated after RNF219 knockdown tended to be changed in the opposite direction after CNOT10 knockdown (Fig 4B). The RNA levels of 42 genes were significantly down-regulated upon RNF219 knockdown, while up-regulated upon CNOT10 knockdown with fold change greater than 1.5 (Fig 4C). For example, the *Zscan4* family genes, which are the specific markers for 2-cell stage embryos and 2-cell state like ES cells, were listed among the 42 RNF219-CNOT10 targets (Fig 4D, S3C) (26). Thus, RNF219 and CNOT10 might play opposite roles in controlling transcript level.

**Figure 4.**
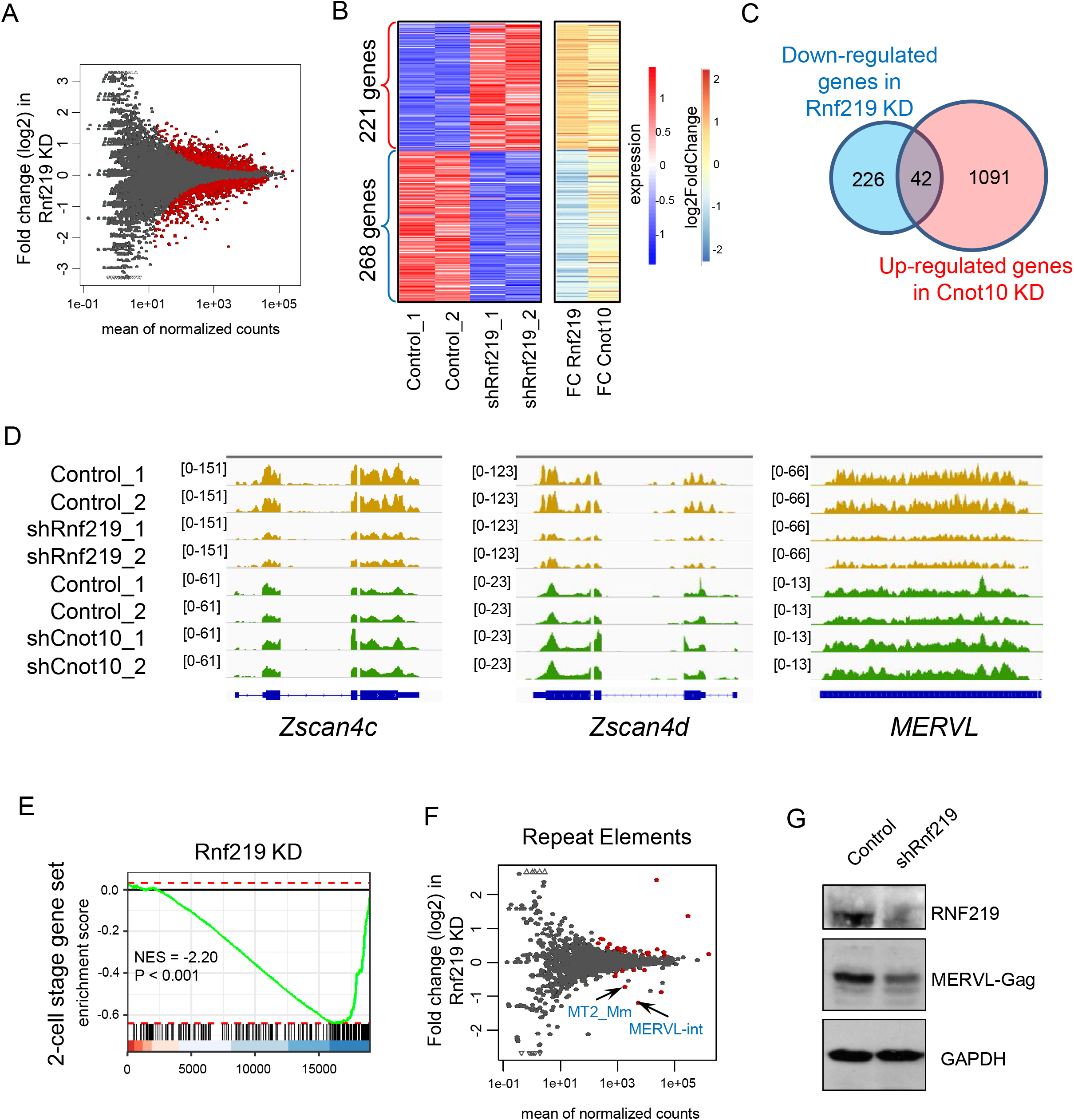
The 2-cell specific genes are oppositely regulated by RNF219 and CNOT10 in mouse ES cells. (A) MA plot showing differential expression of genes after knockdown of RNF219 in mouse ES cells. The MA plot depicts the mean of normalized counts (x-axis) and log2 fold changes that calculated using DESeq2. The red dots represent genes with significance that the adjusted p-value < 0.05. (B) Heat maps showing expression levels (left panel) and fold change (right panel) of differential expressed genes (|Log2FC|>0.58) in RNF219 knockdown mouse ES cells. The right panel also showing fold change of these genes in CNOT10 knockdown mouse ES cells. (C) The Venn diagram depicting the overlap of the down-regulated genes after RNF219 knockdown and the up-regulated genes after CNOT10 KD. |Log2FC|>0.58. (D) Genome browse track file showing that the *Zscan4* family and *MERVL* were down-regulated after RNF219 knockdown, but up-regulated after CNOT10 knockdown. (E) GSEA plot showing that the 2-cell specific genes are highly enriched in the down-regulated genes after RNF219 knockdown in mouse ES cells. (F) MA plot showing differential expression of repeat elements after knockdown of RNF219 in mouse ES cells. The MA plot depicts the mean of normalized counts (x-axis) and log2 fold changes that calculated using DESeq2. The red dots represent repeat elements with significance that the adjusted p-value < 0.05. (G) Western blot showing that the protein level of MERVL-Gag was reduced after RNF219 knockdown in mouse ES cells.

Interestingly, besides *Zscan4c* and *Zscan4d*, 44 of the rest of the genes down-regulated after RNF219 knockdown were 2-cell specific as well. Gene set enrichment analysis (GSEA) confirmed that the 2-cell specific genes were preferentially down-regulated upon RNF219 knockdown (Fig 4E). The *Zscan4* genes are located adjacent to the endogenous retrovirus *MERVL*. Reminiscent of the expression pattern of *Zscan4, MERVL* is also exclusively expressed in 2-cell embryos and 2-cell like ES cells (27). The regulation of 2-C specific gene transcript levels by RNF219 prompted us to further investigate whether *MERVL* transcripts are also targeted by RNF219. We thus analyzed the expression levels of the repeat elements, including *MERVL*, following the RNF219 knockdown by mapping RNA-seq reads to the consensus of different repeat elements. MA-plot analysis indicated that the expression levels of *MERVL* and *MERVL*-derived *LTR MT2 Mm* were also significantly down-regulated following RNF219 knockdown (Fig 4D,F). Consistently, the expression level of MERVL-Gag protein was also reduced in RNF219 knockdown ES cells (Fig 4G).

### Requirement of RNF219 for neuronal specification of mouse ES cells

To investigate whether RNF219 is involved in controlling transcript levels during ES cell differentiation, we performed differential RNA level analyses in retinoid acid (RA) exposed mouse ES cells for neuron differentiation after RNF219 knockdown. RNA-seq analyses identified 400 genes down-regulated and 241 genes up-regulated genes in RA treated mouse ES cells after RNF219 knockdown (Fig 5A,B). The levels of the transcripts from the *Zscan4* family and *MERVL* were also maintained by RNF219 in differentiated ES cells (Fig S5A). Further functional annotation of using DAVID indicated that genes involved in developmental processes and cell differentiation were highly enriched in the down-regulated gene list (Fig 5C). For example, the RNA levels of the known neuronal genes *Neurog1, Neurog3* and *Olig3* were reduced after RNF219 knockdown in RA-treated mouse ES cells (Fig 5D). Interestingly, CNOT10 knockdown upregulated the RNA levels of these genes in RA differentiated cells. To examine the requirement of RNF219 in neuronal specification, a RNF219 knockout (KO) ES cell line was generated and assessed for differentiation ability in in N2B27 media (30) (Fig S5B). The RNF219 KO ES cells displayed typical ES morphology in 2i/LIF culture condition (Fig 5E). However, the cells cannot be maintained and differentiated after 4 days of culture in the N2B27 differentiation media. Our results indicated that RNF219 could be an essential factor for neuron differentiation of ES cells.

**Figure 5.**
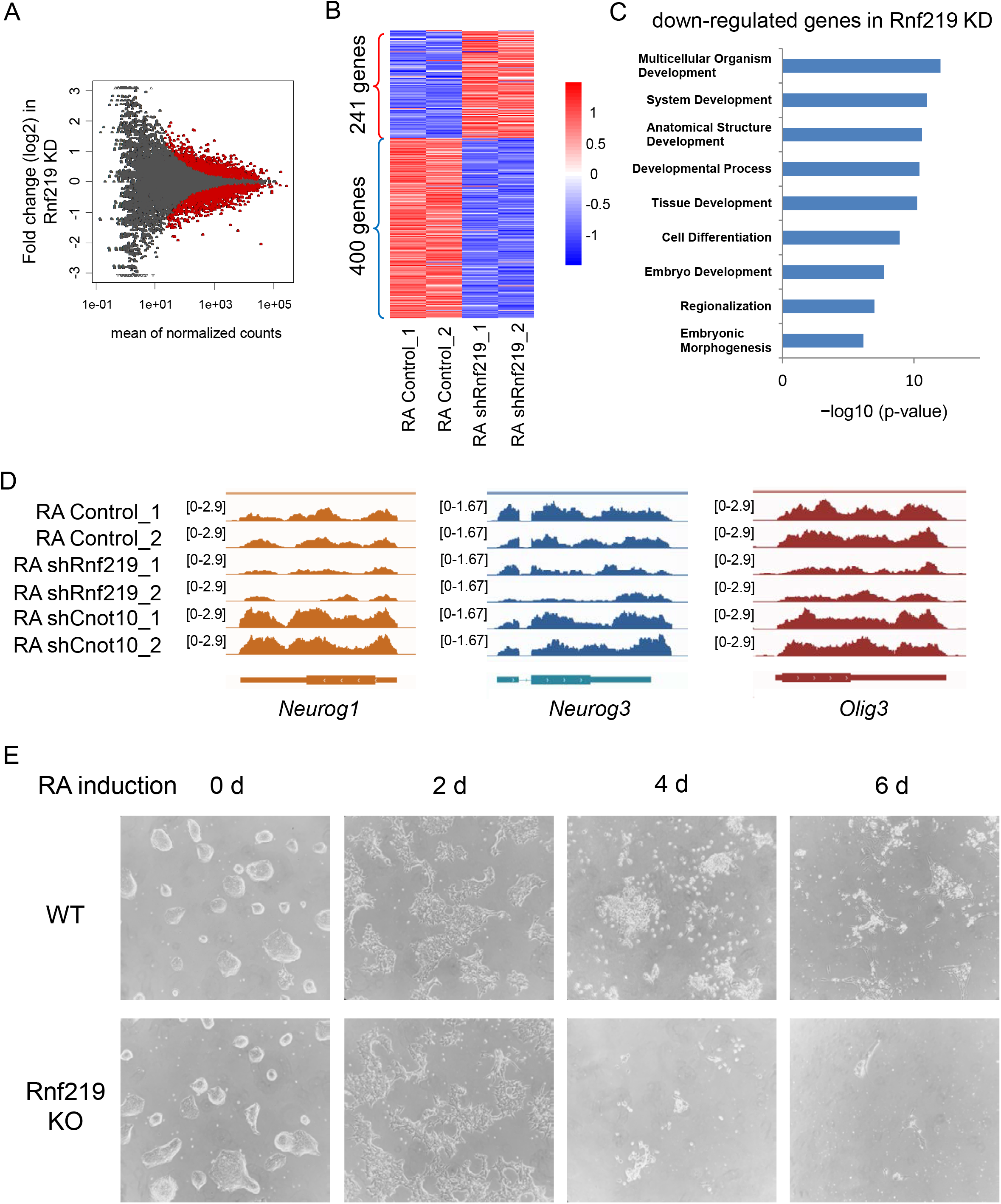
RNF219 is essential for neuronal specification of mouse ES cells. (A) MA plot showing differential expression of genes after RNF219 knockdown in mouse ES cells treated with RA. The MA plot depicts the mean of normalized counts (x-axis) and log2 fold changes that calculated using DESeq2. The red dots represent genes with significance that the adjusted p-value < 0.05. (B) Heat maps showing expression levels of differential expressed genes after RNF219 knockdown in mouse ES cells treated with RA. |Log2FC|>0.58. (C) Functional annotation using DAVID of the down-regulated genes after RNF219 knockdown in mouse ES cells treated with RA. (D) Genome browser track files showing that *Neurog1, Neurog3* and *Olig3* were down-regulated after RNF219 knockdown, but up-regulated after CNOT10 knockdown in mouse ES cells treated with RA. (E) Monolayer differentiation of WT and RNF219 KO mouse ES cells in N2B27 medium for indicated days.

## DISCUSSION

CCR4-NOT is the major deadenylase in mammals. We here demonstrated that RNF219 interacts with CCR4-NOT and functions in ES cell pluripotency maintenance. RNF219 is required for maintaining the proper expression levels of some of the 2-cell specific genes and the neuronal genes in mouse ES cells. RNF219 knockdown impairs the capability of ES cells in neuronal differentiation. Our study thus could highlight a potential direction in investigating the causes of *RNF219* variants related diseases in nervous system.

The interplay between miRISC and CCR4-NOT has been intensively studied. AGO and GW182 are the two core components of miRISC. Following facilitation of miRNA pair with 3’UTR of target RNA by AGO, GW182, or its human paralog TNRC6, recruits CCR4-NOT through interacting with the scaffold CNOT1. In this study, we found that RNF219 can interact with both CCR4-NOT and AGO2, suggesting that a potential double locker mechanism to consolidate miRISC with CCR4-NOT in controlling miRNA mediated deadenylation. Intriguingly, CNOT4, the core subunit of the canonical CCR4-NOT, was not detected in FLAG-RNF219 purification by mass spectrometry. As both CNOT4 and RNF219 are RING domain containing factors possessing E3 ligase activity, we postulated that CNOT4 and RNF219 might be mutually exclusive and that different versions of CCR4-NOT could exist with functional diversity and target specificity.

Opposite regulation by RNF219 and CNOT10 in transcript levels suggests that the deadenylation machinery might be multi-layered with buffering element, which might be of crucial importance to development. For example, *Zscan4* and *MERVL*, whose transcripts are identified in the current study as the *in vivo* targets of RNF219 and CNOT10, are essential for maintenance of telomeres, genome stability and even totipotent state of 2-cell state like ES cells (26,27). Although only sporadic ES cells in culture are ZSCAN4 and MERVL positive at a given time, nearly every ES cell oscillates between ZSCAN4^+^ MERVL^+^ and ZSCAN4^-^ MERVL^-^ states after prolonged passaging (26,27), suggesting that dramatically rapid turnover of the *MERVL* and *Zscan4* transcripts occurs. Slowing down the deadenylation rate of these transcripts by RNF219 might be prone to prevent precocious decay, thus creating a time window enough for MERVL and ZSCAN4 to execute its functions during ES cell maintenance and early embryo development.

## MATERIALS and METHODS

### Antibodies

Antibody against RNF219 was generated in our laboratory. Antibody against CNOT1 was purchased from Proteintech (14276-1-AP). Antibody against CNOT7 was purchased from Abcam (ab195587). Antibody against MERVL-Gag was purchased from Beyotime (AF0240). Antibody against AGO2 was purchased from Millipore (MABE253). Antibodies recognizing TUBULIN, GAPDH, FLAG, V5 or HA were obtained from Sigma.

### Plasmids and cell culture

pV5-CNOT7, pV5-CNOT6, pHA-CNOT1, TBG, BG-L7, BG-PLAC2, BG-1boxB and pNF-YTHDF2 were previously described (18). FLAG-tagged RNF219 and CNOT1 cDNAs were cloned into pCDNA5/FRT-TO vector (Invitrogen). The expression vectors were then transfected into HEK-293 Flp-In-TRex cells and followed by selection with hygromycin to generate stable cell lines. HeLa-tTA cells were purchased from Clontech. V6.5 mouse ES cells were cultured in serum and LIF supplemented medium on irradiated mouse embryonic fibroblasts (MEFs). All cells were maintained at 37°C under 5% CO2.

Lentivirus mediated RNAi was previously described (31). 72 hours after lentiviral infection, ES cells were treated with or without RA for 24 hours before harvesting. For all analyses, cells were grown for one passage off feeders on tissue culture plates for 30 min. RNF219 knockout V6.5 ES cell line was generated by CRISPR-Cas9 genome editing technique. For neuron differentiation, mouse ES cells were cultured in N2B27 medium and treated with RA for 6 days.

### Flag purification, immunoprecipitation, and size exclusion chromatography

Flag purification was performed as previously described (32,33). Briefly, the expression of FLAG-RNF219 was induced with doxycycline for 48 hours. The protein complexes were purified using the FLAG-affinity purification approach in the presence of benzonase (Sigma), and analyzed by SDS-PAGE and silver staining before subjected to Mass Spectrometry.

Immunoprecipitation was performed as previously described (32,33). Briefly, cells were lysed in 420 mM NaCl containing lysis buffer supplemented with protease inhibitor cocktail (Sigma) at 4°C. After centrifugation, the balance buffer (20 mM HEPES [pH 7.4], 1 mM MgCl2, 10 mM KCl) was added to the supernatant to make the final NaCl concentration 300 mM. The supernatant was then incubated with antibodies and protein A beads overnight at 4°C. The beads were spun down and washed three times with the wash buffer before boiling in the SDS loading buffer.

Size exclusion chromatography was performed as previously described (32,33). Nuclear and cytoplasmic extracts were subjected to Superose 6 size exclusion chromatography (GE Healthcare) with size exclusion buffer (40 mM HEPES [PH 7.5], 350 mM NaCl, 10% glycerol and 0.1% Tween-20). Fractions were resolved in SDS-PAGE gels, followed by Western blotting.

### *In vitro* deadenylation assay

HEK-293 FIT cells stably expressing FLAG-CNOT1 or FLAG-RNF219 were subjected to FLAG purification. The FLAG-M2 beads bound bait protein and it interacting factors were incubated with 200 nM FAM-N20A20 synthetic RNA substrate in 50 mM Tris-HCl [pH 7.5], 50 mM NaCl, 2 mM MgCl2 and 0.5 mM DTT in a rotator at 37°C for 30 min. Samples were collected at different time points and analyzed on a 7 M urea 20% polyacrylamide denaturing gel. The images were analyzed with a fluorescence imager.

### Reporter deadenylation assay

Deadenylation assay of BG reporters was previously described (18) with pcDNA5-RNF219 or truncations overexpressed. HeLa-tTA cells were plated on 6-well plate 1 day before transfection in DMEM containing 20 ng/ml tetracycline. 1 ug of the reporter plasmid and 500 ng of RNF219 or control plasmid were used for transfection. The transcription of BG mRNA was induced by removing tetracycline 12 hours after transfection. 3 hours after induction, tetracycline was added to a final concentration of 1 mg/ml to block the transcription of BG. Cytoplasmic RNAs were then isolated at various time intervals. RNAs were treated with RNaseH in the presence of an antisense DNA oligo (5’-GTCCAGGTGACTCAGACCCTC-3’ for TBG, BG-L7 and BG-1boxB; 5’-CCAGCCACCACCTTCTGATAGGC-3’ for BG-PLAC2). The digested RNA samples were then analyzed by electrophoresis (5.5% PAGE with 8 M urea) and northern blotting using the DIG Northern Starter Kit (Roche). Poly(A) tail assay of endogenous ZSCAN4C was measured by Poly(A) Tail-Length Assay Kit (Thermo, 764551KT).

### RNA-seq data processing

We first evaluated each RNA-seq data quality using FastQC (version 0.11.8) and confirmed that all datasets are qualified for following analysis. The RNA-seq reads were aligned to mouse reference genome (mm10/GRCm38) using HISAT2 (version 2.1.0). The mouse reference genome sequence was downloaded from ENSEMBL (Mus Musculus GRCm38/mm10). Then we used featureCounts (version 2.0.0) to count reads or read pairs for each protein-coding gene annotated in GENCODE (vM23)(34). Besides, we also counted reads or read pairs for repeat elements annotated in RepeatMasker. Then we conducted differential expression analysis using DESeq2(35) and selected the |log2FoldChange|> 1.5 and FDR < 0.05 as the cutoff to identify significantly differential expressed genes.

### Identification of 2-cell stage genes

To identify those genes that were specifically expressed in mouse 2-cell stage, we downloaded a single-cell RNA-seq dataset (GSE53386) from GEO(36). The gene expression profiles (FPKM) of oocyte, zygote, 2-cell, 4-cell, 8-cell. Blastocyst and morula were collected. Then we calculated a specificity ratio for each gene and we selected specificity ratio > 1 as the cutoff to identity cell type specific genes for each cell stage.

### Function enrichment analysis

For GO enrichment analysis, we used g:profiler to estimate enrichment for each gene set(37). EnrichmentMap was used to visualize the enrichment results(38).

For GSEA analysis, we used the R package fgsea to calculate the enrichment score and p-value. The input gene sets file (.gmt) were downloaded from g:profiler. The enrichment results were visualized using R.

### RNA stability analysis

We used REMBRANDTS to estimate differential mRNA stability using RNA-seq data across multiple samples. Δexon–Δintron was used as an estimate of differential mRNA stability in several recent studies(39,40). REMBRANDTS is designed to estimate gene-specific bias function that is then subtracted from Δexon–Δintron to provide unbiased differential mRNA stability measure(41). The coordinates of exonic and intronic segments were extracted from GENCODE (vM23). HTSeq was used to count reads that map to exonic or intronic segments for each gene.

## Acknowledgments

The authors are grateful to the members in Lin Lab for critical discussion, and Toby Wai Kiat Chin for technical assistance of this study.

## SUPPLEMENTART FIGURE LEGENDS

**Figure S1.**
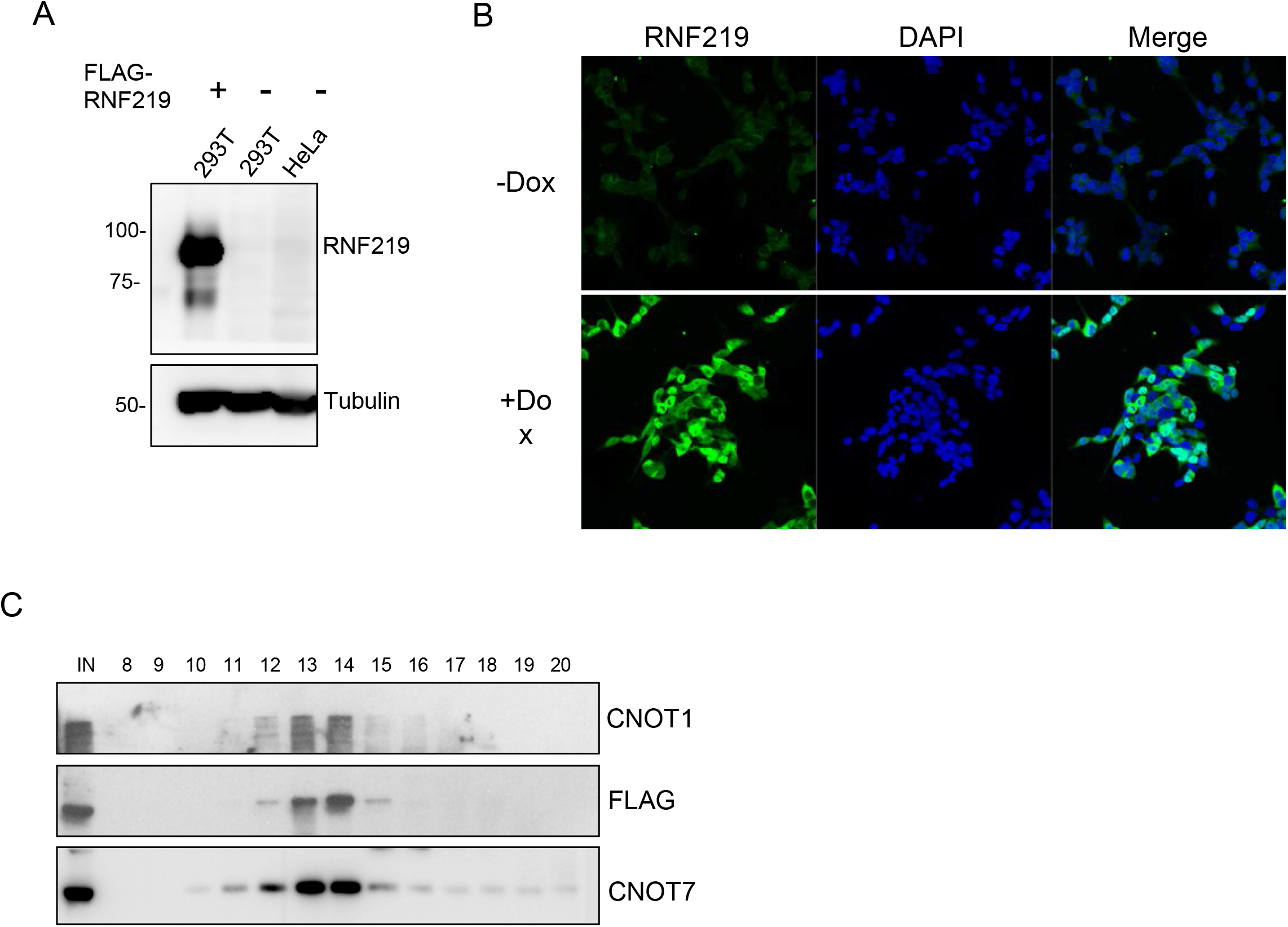
Interaction of RNF219 with the components of CCR4-NOT. (A) Western blot showing that the RNF219 antibody used in this study was able to recognize ectopically expressed FLAG-RNF219 in HEK-293 cells. (B) Immunofluorescence showing that the RNF219 antibody was able to recognize ectopically expressed FLAG-RNF219. (C) Size exclusion chromatography of nuclear extracts from the FLAG-RNF219 stable cell line demonstrating that majority of RNF219 co-eluted with the components of CCR4-NOT.

**Figure S2.**
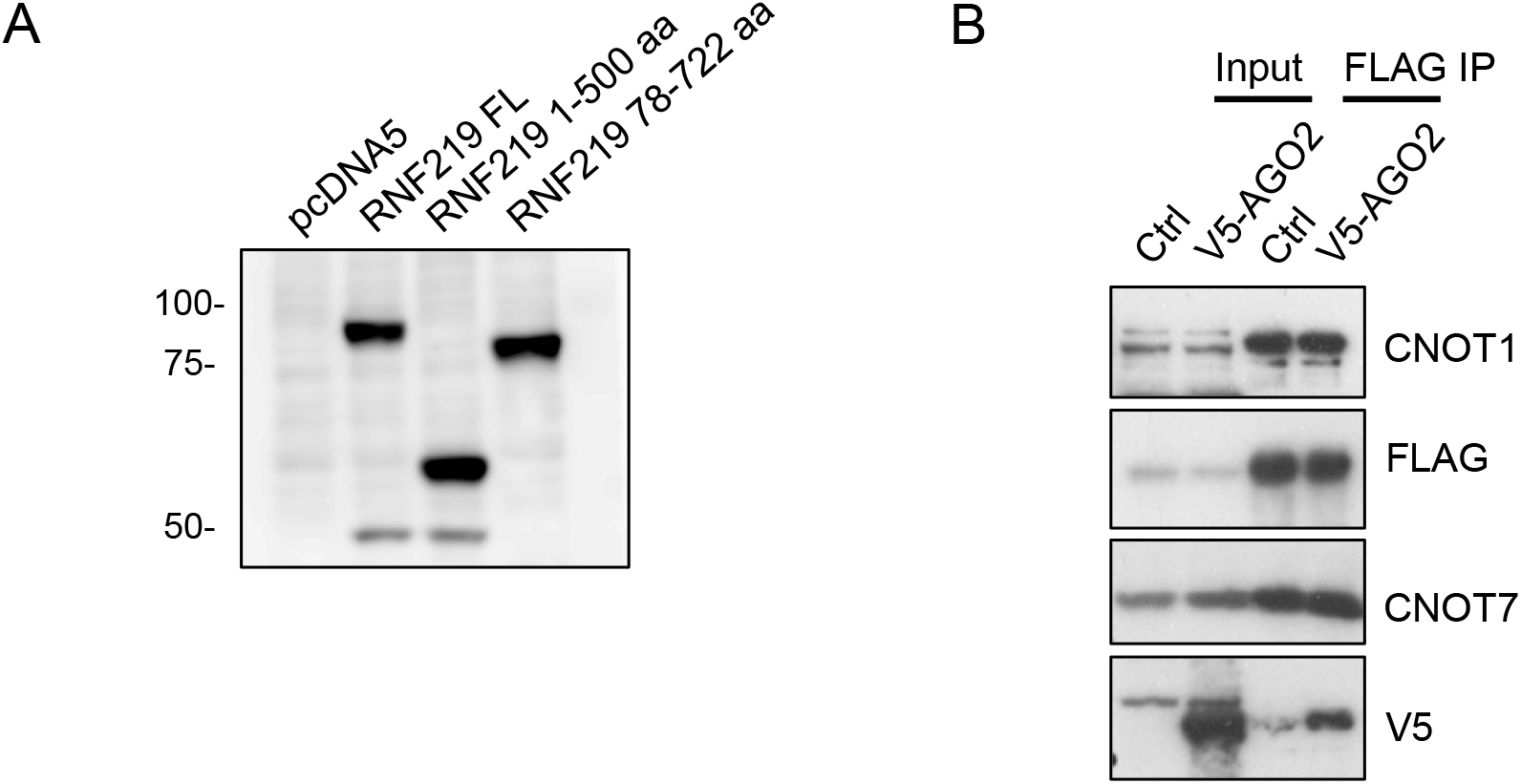
Interaction of RNF219 with AGO2. (A) Western blot showing overexpression of RNF219 full length, 1-500 aa and 78-722 aa truncation proteins used in deadenylation assays. (B) Interaction of RNF219 with AGO2. V5-AGO2 was expressed in FLAG-RNF219 stable cell lines. The cells were then subjected to lysis, and FLAG immunoprecipitation.

**Figure S3.**
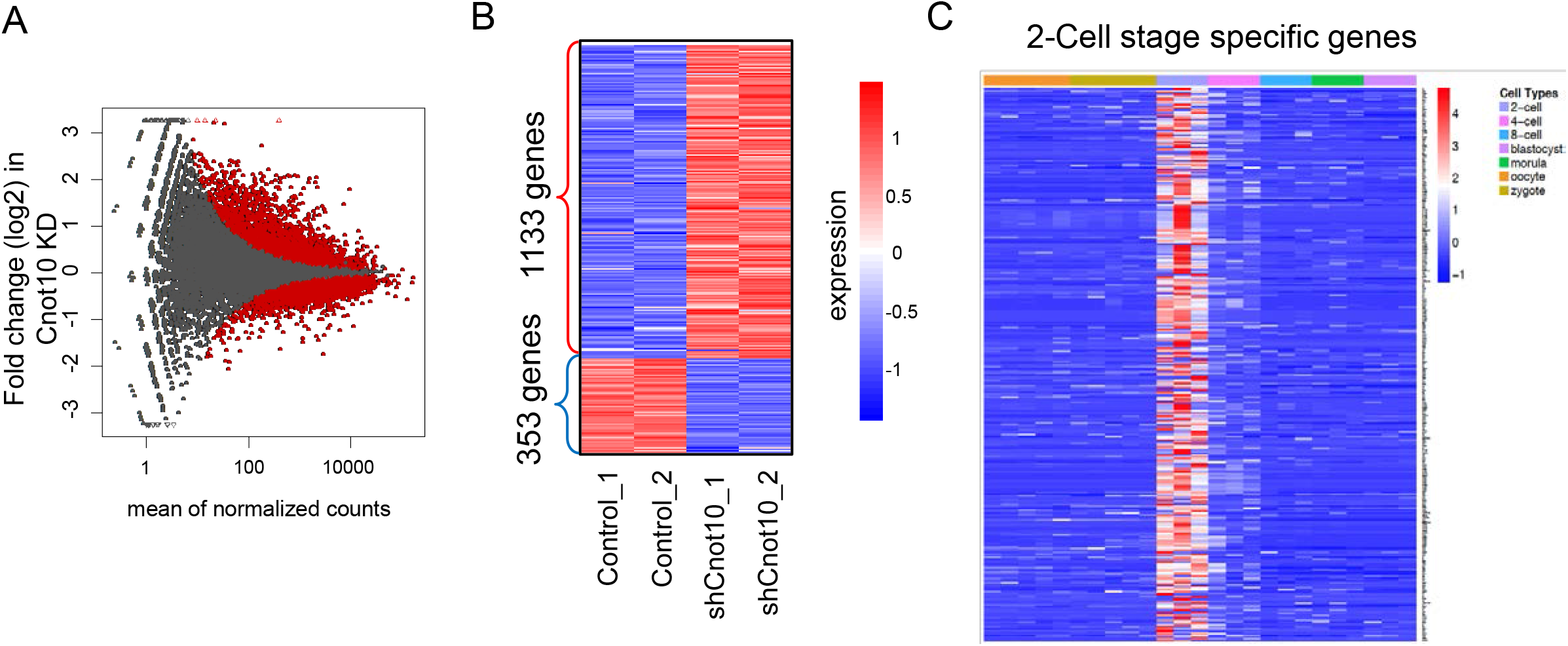
Genome wide analyses of gene expression change after CNOT10 knockdown. (A) MA plot showing differential expression of genes after knockdown of CNOT10 in mouse ES cells. The MA plot depicts the mean of normalized counts (x-axis) and log2 fold changes that calculated using DESeq2. The red dots represent genes with significance that the adjusted p-value < 0.05. (B) Heat maps showing expression levels of differential expressed genes (|Log2FC|>0.58) in CNOT10 knockdown mouse ES cells. (C) Heat maps showing the specificity ratio for each 2-cell stage genes in each cell type.

**Figure S4.**
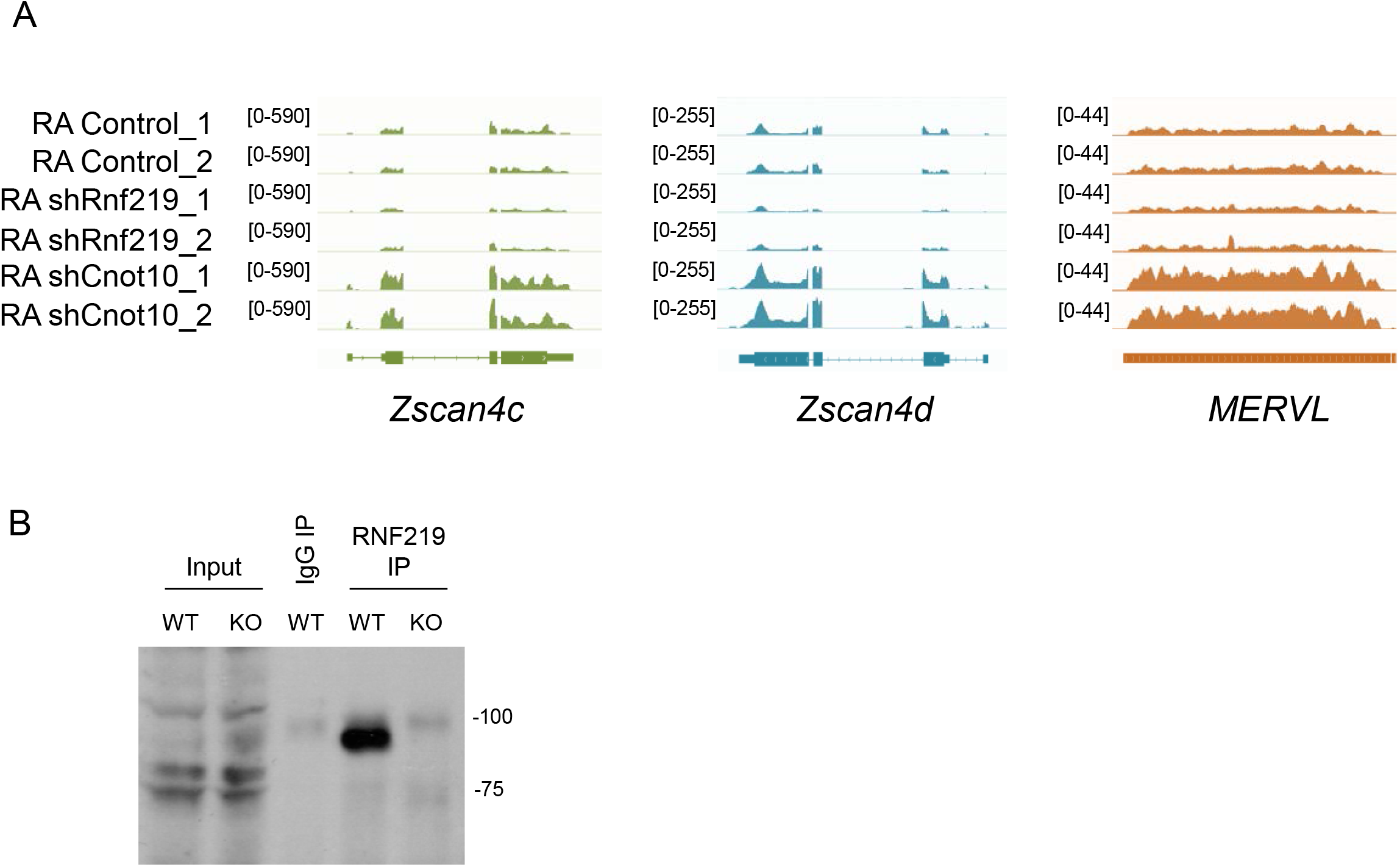
Oppositely regulation of *Zscan4* and *MERVL* by RNF219 and CNOT10 in mouse ES cells treated with RA. (A) Genome browse track file showing that the *Zscan4* family and *MERVL* were down-regulated after RNF219 knockdown, but up-regulated after CNOT10 knockdown in mouse ES cells treated with RA. (B) Validation of the RNF219 KO ES cell line generated in this study by immunoprecipitation, followed by western blot.

